# Identification of beneficial and detrimental bacteria that impact sorghum responses to drought using multi-scale and multi-system microbiome comparisons

**DOI:** 10.1101/2021.04.13.437608

**Authors:** Mingsheng Qi, Jeffrey C. Berry, Kira Veley, Lily O’Connor, Omri M. Finkel, Isai Salas-González, Molly Kuhs, Julietta Jupe, Emily Holcomb, Tijana Glavina del Rio, Cody Creech, Peng Liu, Susannah Tringe, Jeffery L. Dangl, Daniel Schachtman, Rebecca S. Bart

**Affiliations:** Donald Danforth Plant Science Center, St. Louis, MO, USA; Department of Biology, University of North Carolina at Chapel Hill, Chapel Hill, NC, USA; Howard Hughes Medical Institute, University of North Carolina at Chapel Hill, Chapel Hill, NC, USA; Department of Plant and Environmental Sciences, Institute of Life Science, The Hebrew University of Jerusalem, Jerusalem, Israel; Curriculum in Bioinformatics and Computational Biology, University of North Carolina at Chapel Hill, Chapel Hill, NC, USA; DOE Joint Genome Institute, Lawrence Berkeley National Laboratory, Berkeley, CA, USA; Environmental Genomics and Systems Biology Division, Lawrence Berkeley National Laboratory, Berkeley, CA, USA; Department of Agronomy and Horticulture, University of Nebraska-Lincoln, Scottsbluff, NE, USA; Department of Statistics, Iowa State University, Ames, IA, USA; Carolina Center for Genome Sciences, University of North Carolina at Chapel Hill, Chapel Hill, NC, USA; Curriculum in Genetics and Molecular Biology, University of North Carolina at Chapel Hill, Chapel Hill, NC, USA; Department of Microbiology and Immunology, University of North Carolina at Chapel Hill, Chapel Hill, NC, USA; Center for Plant Science Innovation, University of Nebraska – Lincoln, Lincoln, NE, USA

**Keywords:** Root microbiome, sorghum, drought stress, *Variovorax*, *Arthrobacter*

## Abstract

**Background:** Drought is a major abiotic stress that limits agricultural productivity. Previous field-level experiments have demonstrated that drought decreases microbiome diversity in the root and rhizosphere and may lead to enrichment of specific groups of microbes, such as *Actinobacteria*. How these changes ultimately affect plant health is not well understood. In parallel, model systems have been used to tease apart the specific interactions between plants and single, or small groups of microbes. However, translating this work into crop species and achieving increased crop yields within noisy field settings remains a challenge. Thus, the next scientific leap forward in microbiome research must cross the great lab-to-field divide. Toward this end, we combined reductionist, transitional and ecological approaches, applied to the staple cereal crop sorghum to identify key beneficial and detrimental, root associated microbes that robustly affect drought stressed plant phenotypes.

**Results:** Fifty-three bacterial strains, originally characterized for association with *Arabidopsis*, were applied to sorghum seeds and their effect on root growth was monitored for seven days. Two *Arthrobacter* strains, members of the *Actinobacteria* phylum, caused root growth inhibition (RGI) in *Arabidopsis* and sorghum. In the context of synthetic communities, strains of *Variovorax* were able to protect both *Arabidopsis* and sorghum from the RGI caused by *Arthrobacter*. As a transitional system, we tested the synthetic communities through a 24-day high-throughput sorghum phenotyping assay and found that during drought stress, plants colonized by *Arthrobacter* were significantly smaller and had reduced leaf water content as compared to control plants. However, plants colonized by both *Arthrobacter* and *Variovorax* performed as well or better than control plants. In parallel, we performed a field trial wherein sorghum was evaluated across well-watered and drought conditions. Drought responsive microbes were identified, including an enrichment in *Actinobacteria*, consistent with previous findings. By incorporating data on soil properties into the microbiome analysis, we accounted for experimental noise with a newly developed method and were then able to observe that the abundance of *Arthrobacter* strains negatively correlated with plant growth. Having validated this approach, we cross-referenced datasets from the high-throughput phenotyping and field experiments and report a list of high confidence bacterial taxa that positively associated with plant growth under drought stress.

**Conclusions:** A three-tiered experimental system connected reductionist and ecological approaches and identified beneficial and deleterious bacterial strains for sorghum under drought stress.

## Introduction

Many factors influence overall plant health and productivity including varietal differences (*G_P_*), abiotic stresses (*E*) and the diverse collection of microbes (*G_M_*) that live intimately in and around plants (Buée et al. 2009; Rolli et al. 2015; Agler et al. 2016). The composition as well as the spatial and temporal dynamics of the plant microbiota are also influenced by environmental conditions and host factors (Bogino et al. 2013; Bodenhausen et al. 2014; Bouffaud et al. 2014; Naylor et al. 2017; Samad et al. 2017; Fitzpatrick et al. 2018; Xu et al. 2018) resulting in a tangled web of interactions (*PtantHealth* = *G_P_* × *G_M_* × *E*). Previous research aimed at untangling this web of interactions can be divided into two general approaches: field-based surveys and controlled system experiments.

In field-based surveys, next generation amplicon sequencing is used to directly quantify microbial constituents associated with plants, often across various abiotic stresses (Lauber et al. 2008; Rousk et al. 2010; Xu et al. 2018). These experiments group microbes into taxonomic units, often at the family or genus level, and are useful for observing major community shifts/differences. For example, it is well documented that compared to bulk soil, the root and rhizosphere contain much less microbial diversity, suggesting that plant roots influence the composition of their microbiomes (Edwards et al. 2015; Naylor et al. 2017; Fitzpatrick et al. 2018). Similarly, previous studies have shown that drought decreases the diversity of microbes in the roots of 30 angiosperm plants and 18 grass crop species including sorghum (Naylor et al. 2017; Fitzpatrick et al. 2018; Xu et al. 2018). Notably, in these studies, *Actinobacteria* strains were enriched in both bulk soil and even more enriched in roots. From this observation, it was hypothesized that these gram-positive (monoderm) bacteria display inherent physiological adaptation to drought as well as a response to plant metabolic changes under drought. Furthermore, some studies suggest that when plant hosts suffer from abiotic/biotic stresses, they recruit specific microbes able to alleviate the stress, known as the ‘‘cry for help” hypothesis (Berendsen et al. 2018; Harbort et al. 2020).

Biological nitrogen fixation and improved nutrient uptake by the mutualistic symbioses between legumes and rhizobia and between cereals and mycorrhizae, respectively, are among the most well-characterized examples of plant growth-promoting microbial processes and have been successfully studied in both lab and field settings (Sessitsch and Mitter 2015). Reductionist experiments within controlled systems have been used to probe the specific function of many additional beneficial microbes. However, in general, translating microbe-derived plant growth promoting phenotypes from labs into complex agricultural settings remains a challenge. For example, while *Azospirillum brasilense* strains promoted vegetative growth of maize and wheat in the greenhouse, they had little impact on plant growth in the field (Fukami et al. 2016). Recent efforts have used microbial synthetic communities (SynComs) as a reductionist model for natural microbiota. SynComs have been used to decipher *in planta* processes that lead to plant-microbiota homeostasis and to understand the mechanisms underlying the microbiota’s effects on plant growth, nutrient uptake and disease resistance (Berendsen et al. 2018; Herrera Paredes et al. 2018; Voges et al. 2019; Finkel et al. 2020). Berendsen et al discovered three rhizosphere bacterial species that are specifically enriched upon *Arabidopsis* foliar defense activation by the downy mildew pathogen (Berendsen et al. 2018). These three strains were able to function synergistically in the field soils and induce systemic resistance to downy mildew disease. Voges et al observed that iron deficiency caused a compositional shift to a SynCom in *Arabidopsis* roots and this was linked to changes in root exudation (Voges et al. 2019). Most recently, using top-down deconstruction of a large phylogenetically diverse SynCom, it was demonstrated that the bacterial genus *Variovorax* (Finkel et al. 2020), a core rhizosphere member across plant species and geographic locations (Fitzpatrick et al. 2018; Finkel et al. 2020; Thiergart et al. 2020), was able to protect *Arabidopsis* root growth from diverse root growth inhibitory strains. *Variovorax* strains protect the host plant from manipulation by hormone-secreting microbes within the microbiome, suggesting chemical interference as a novel strategy that enhances plant resilience.

Drought is one of the most important abiotic stresses for crop plants and sorghum [*Sorghum bicolor* (L.) Moench] is one of the best-adapted cereal crops to water-limited environments (Ludlow and Muchow 1990). Decades of breeding have resulted in elite sorghum varieties and hybrids with optimized drought tolerance traits (*G_P_* × *E*) including waxy leaf surfaces, deep root systems and the ‘stay green’ trait (Fracasso, Trindade, and Amaducci 2016; Kamal et al. 2019). In contrast, interactions between the root-associated microbiome and drought (*G_M_* × *E*) and between the plant and the root-associated microbiome (*G_P_* × *G_M_*), are less well understood.

Here, we test bacterial SynComs that affects *Arabidopsis* root growth (Finkel et al. 2020) to determine whether a similar microbe-dependent phenotype is observed on sorghum. We tested the SynComs in a sorghum germination assay and a sorghum phenotyping assay and found that *Arabidopsis*-protective *Variovorax* strains can also protect sorghum growth from drought and root growth inhibition (RGI) from various bacterial strains. In parallel, we performed a sorghum field trial with well-watered and drought conditions. Drought-responsive microbes were identified including an enrichment of *Actinobacteria*, consistent with previous findings. Additionally, sorghum-associated bacteria, both beneficial and deleterious, were discovered from the phenotyping assay and the field trial. Several bacteria were observed to have phenotypic effects in both systems and so become high-priority candidates for future study. All three datasets suggest that *Arthrobacter* strains impair sorghum growth, especially under drought stress. To our knowledge, this is the first example of reductionist and ecological approaches revealing convergent results on crop plant associated microbial interactions relevant for a specific host plant trait.

## Materials and Methods

### Sorghum root growth assay using germination paper. (Fig.1, Fig. S1)

**Fig. 1.**
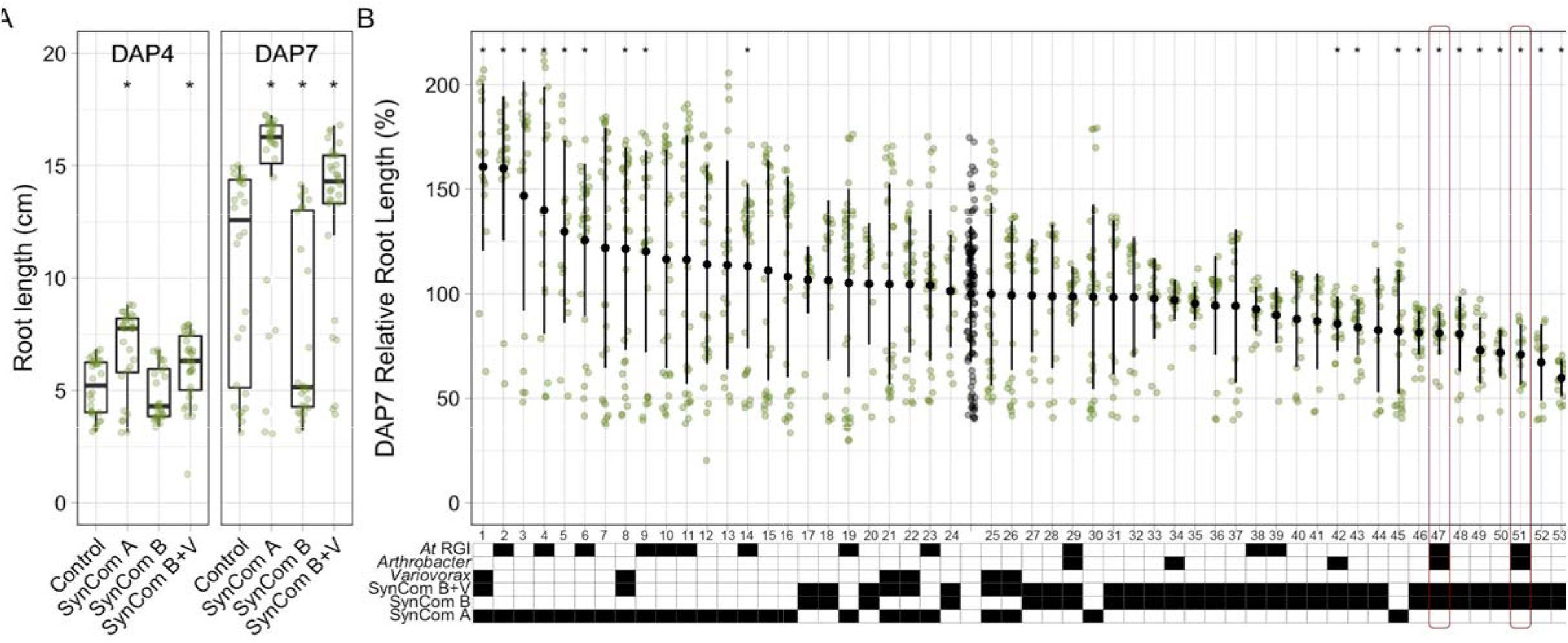
Root length phenotypes after inoculation of sorghum seeds with SynCom strains, as communities (A) or individually (B). Green dots represent the root lengths of individual sorghum seedlings. A. Box plots display medians (horizontal line) the 75th and 25th percentiles (top and bottom of box) and the upper and lower whiskers extend to data no more than 1.5× the interquartile range from the upper edge and lower edge of the box, respectively. B. Each strain was tested individually (1-53) for effect on sorghum seedling root growth. Additional strain details can be found in Supplemental Table 1. Grey dots represent the control (no bacterial treatment) seedlings. The solid black dots and lines represent the mean ± standard deviation. Specific features of each strain are summarized in the table. Black shading indicates that the strain has that feature. RGI: Root Growth Inhibition. Red outline indicates *Arthrobacter* strains (47 and 51) that cause RGI in both *Arabidopsis* and sorghum. The number of replicated samples for each treatment (A and B) n > 20. Wilcoxon tests were performed between SynCom treatments and control samples (sorghum without microbial treatments) (A and B) and p values were corrected using the method of Benjamini-Hochberg. *, p < 0.05.

#### Bacterial cultures

Detailed description of the 53 bacterial strains used in this work have been published elsewhere (Levy et al. 2017; Herrera Paredes et al. 2018; Finkel et al. 2020). Six days before each experiment, bacteria were streaked on NYGA plates with cycloheximide (5 g/L bactopeptone, 3 g/L yeast extract, and 20 mL/L glycerol, with 15 g/L agar for solid medium, 100 mg/L cycloheximide) from glycerol stocks. Bacteria were grown at 30 °C. After 4 days of growth, bacterial strains were re-streaked on fresh NYGA plates with cycloheximide and returned to the incubator for an additional 48 h of growth. Bacteria were resuspended into autoclaved, distilled water (optical density at 600 nm (OD_600_) = 0.5). For the synthetic communities (SynComs), equal volumes of individual bacterial cultures (OD_600_ = 0.5) were combined such that the final inoculum was OD_600_ = 0.5.

#### Plant inoculation, growth, imaging and analysis

*Sorghum bicolor* (L.) Moench BTx623 seeds were surface sterilized in an airtight desiccator with chlorine gas by mixing 40 ml bleach with 5 ml saturated hydrochloric acid for 3 h and then were soaked in 10 ml of sterile water (Control) or bacterial inoculum (individual strains or SynComs), overnight at room temperature. The soaked seeds were placed in the seed pockets of the germination pouch (CYG-38LB, PhytoAB, San Jose, CA). The germination pouches were placed vertically in dark folders, hung in file crates and plants were grown under a 14-h light/10-h dark regime (except all dark for the first day) with temperatures of 30-°C day/25-°C night and 50% humidity. Sorghum roots were imaged four and seven days after planting (DAP), using a document scanner. Primary root length elongation was manually measured using ImageJ. Primary root lengths were compared using a Wilcoxon test. P-values were corrected using the method of Benjamini-Hochberg to control FDR. Significance of differences between treatments were indicated with the asterisks showing adjusted p-value levels.

#### Quantification of the colonized bacteria

For the rhizosphere versus endosphere assays (Fig S1), seeds were treated as above except that bacteria were grown directly from glycerol stocks for 48 hours on plates and then seeds were inoculated with a 1.5×10^8^ CFU/ml (OD_600_ = 0.031 for *Variovorax* and OD_600_ = 0.3 for *Arthrobacter*). Seven DAP, two centimeter long root tip sections from two seedlings were cut and resuspended in 200 ul of wash buffer (10 mM of MgCl_2_, 0.05% Silvet-L77) by vortexing at maximum speed for 5 min. The supernatant, representing the rhizosphere, was transferred to a new tube and bacterial populations were determined by counting colony forming units (CFUs) of serial dilutions. The root sections were then surface-sterilized with a bleach solution (1% bleach, 0.1% Triton-X 100) for 4 min, followed by one wash with 70% ethanol and three washes with wash buffer. Aliquots of the final washes were plated on NYGA plates to determine effectiveness of root surface sterilization. The surface sterilized root sections were transferred into clean 2-ml Safe-lock tubes (Eppendorf) with stainless steel beads and 200 ul of wash buffer, and were homogenized with a TissueLyser II (Qiagen) at 30 Hz for 1 min. The crushed root tissue solutions, representing endosphere, were used for serial dilutions. Aliquots of the dilutions were spread on NYGA plates. The resulting colonies were counted after 2-day incubation at 30°C and calculated as CFU per unit of plant root tissue.

### Sorghum growth assay using the Lemnatec High-throughput phenotyping platform (Fig. 2–3, Fig. S2–S4)

**Fig. 2.**
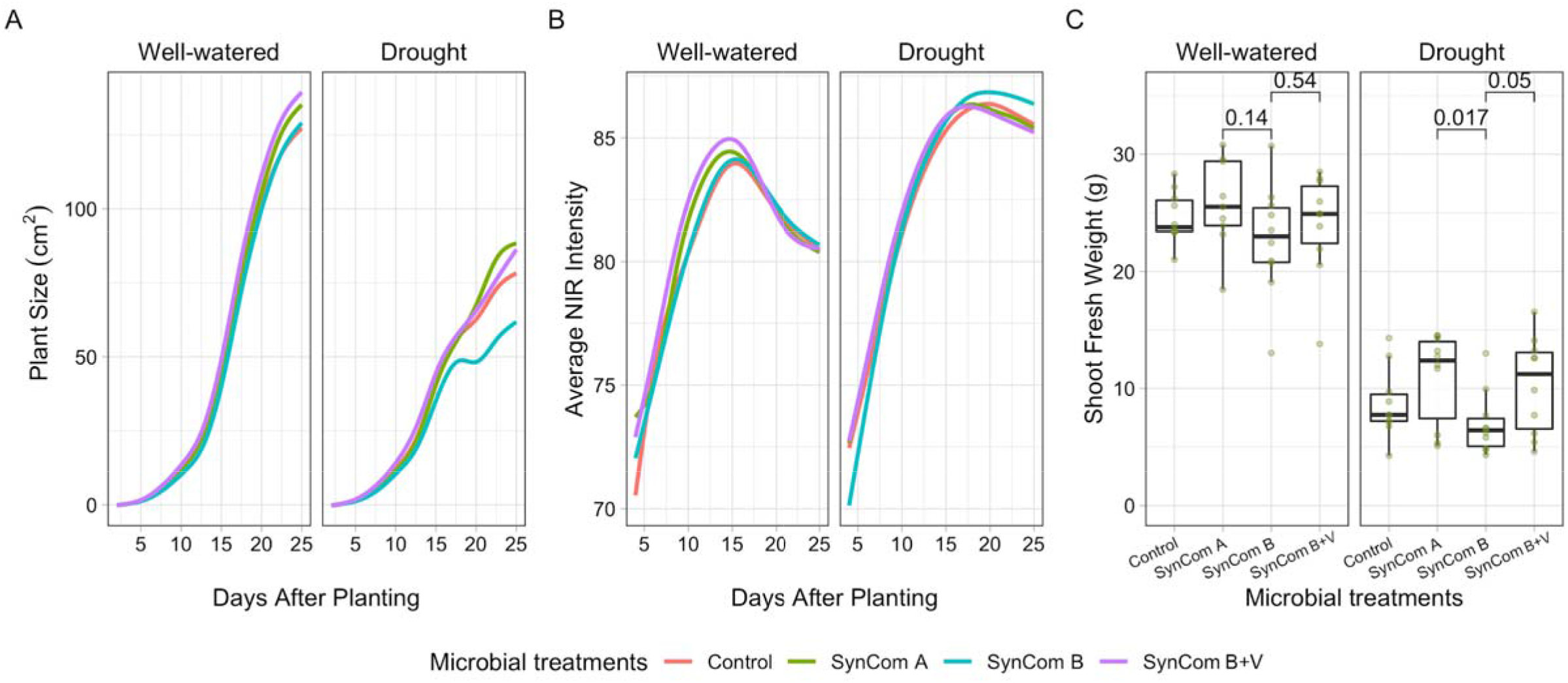
Sorghum growth phenotypes after seed inoculation with SynCom strains quantified in a high-throughput phenotyping assay. The temporal changes of plant size (A) and NIR signal (B) were plotted, with line colors showing the microbial treatments. C. The green dots represent the shoot fresh weight of sorghum at the conclusion of the assay. Box plots display medians (horizontal line) the 75th and 25th percentiles (top and bottom of box) and the upper and lower whiskers extend to data no more than 1.5× the interquartile range from the upper edge and lower edge of the box, respectively. Pairwise t-tests were performed between microbial treatments for well-watered and drought conditions. P-values for select comparisons are shown and all others were not significant at the alpha of 0.05. The number of replicated samples for each treatment n = 50 (A and B) or n = 10 (C).

**Fig. 3.**
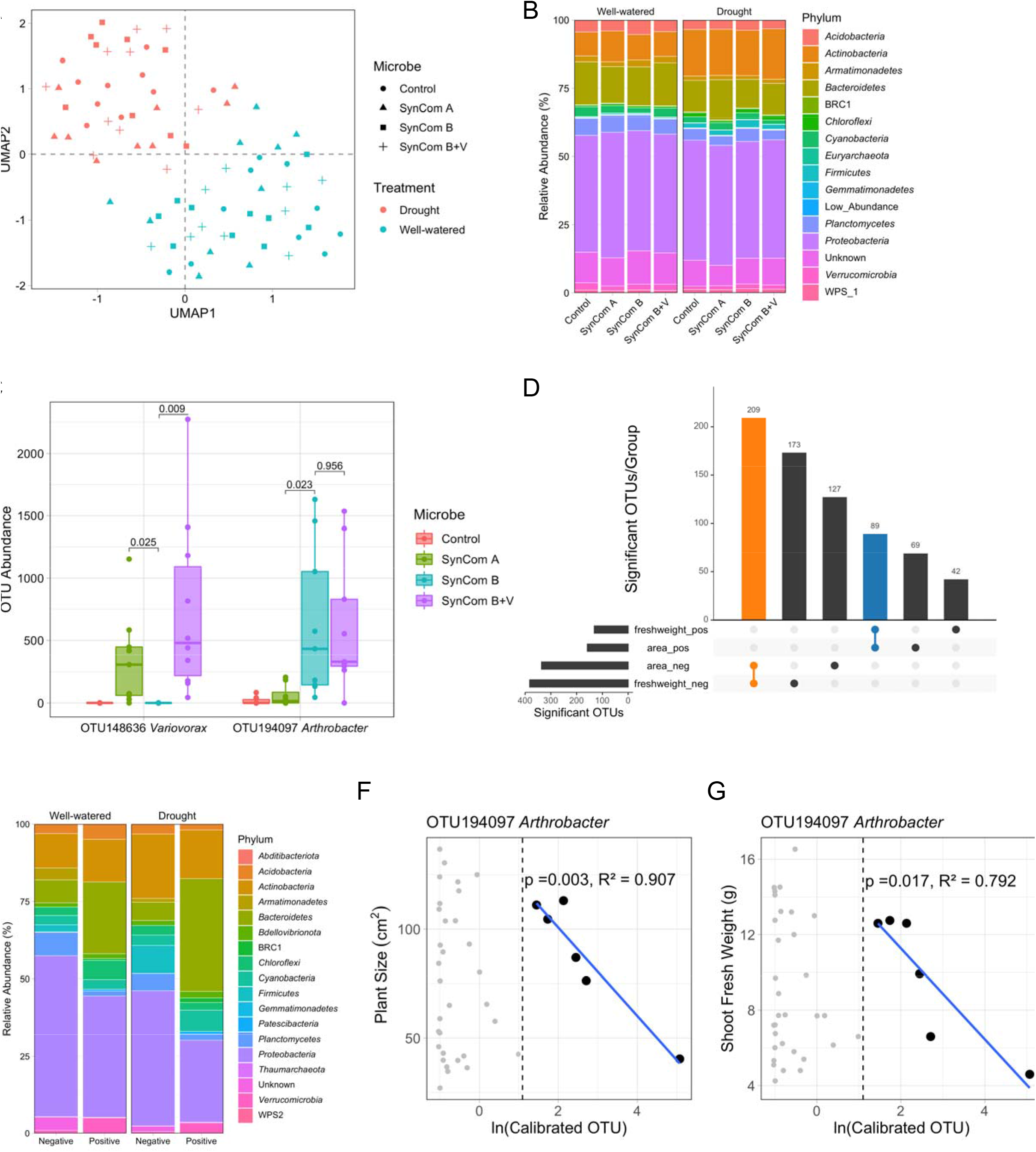
The sorghum root-associated microbiome with SynCom and drought treatments in the high-throughput phenotyping assay. A. The clustering of microbiome samples using unsupervised UMAP, with colors and shapes showing the drought and microbial treatments, respectively. B. Phylum-level distribution of the microbial treatmentspecific microbiota within drought treatments. C. The OTU abundance of *Variovorax* and *Arthrobacter* OTUs at the conclusion of the assay under drought. The dots represent the OTU abundance in different samples with colors showing the microbial treatments. The horizontal bars within boxes represent medians. The tops and bottoms of the boxes represent the 75th and 25th percentiles, respectively. The upper and lower whiskers extend to data no more than 1.5× the interquartile range from the upper edge and lower edge of the box, respectively. Pairwise t-tests were performed between SynCom treatments. P-values for select comparisons are shown and all others were not significant at the alpha of 0.05. The numbers of replicated samples n = 8. D. The numbers of OTUs associated to both plant phenotypes. Colors represent the OTU groups with same association directions. E. Phylum-level distribution of the plant phenotype-associated microbiota within drought treatments. F, G. The changepoint model fitting between OTU abundance and plant phenotypes (F, plant size; G, shoot fresh weight) for OTU194097 *Arthrobacter* strain. Grey dots indicate samples that did not meet the abundance threshold.

#### Bacterial culture and plant inoculation

SynCom strains were prepared for inoculation as described above. Each surface-sterilized sorghum seed was sown 2 inches deep into autoclaved foam plugs (Oasis, Kent, OH) (2 inch deep). Each SynCom inoculum was adjusted to OD_600_ = 0.5 and 13 ml of the corresponding microbial inoculants was poured over the foam plug.

#### Lemnatec plant growth conditions

Sorghum seeds, with the microbial inoculants, were germinated in a Conviron growth chamber set to a 16hr day cycle with temperatures of 32/22 °C and humidity of 60%/40% at day and night, respectively. After 2 days growth, germinated plugs were then transplanted into pre-filled, steam-sterilized, small tree pots (3X3X8 in.) with a one-to-one blend of Metro mix 360 and turface (Hummert International, Earth City, Missouri), that was water-saturated prior to transplanting. Each pot was loaded onto the Bellwether Phenotyping Platform. Growth conditions on the platform were set on a 16 hr cycle with temperatures of 32/22 °C and humidity of 60%/40% at day and night, respectively. Lighting was supplied by metal halide and high-pressure sodium bulbs set to emit 400 *μ*mol*m^−2^*s^−1^. Water was delivered to the plants once per day by adding water to match the target weight of the given treatment. Target weights of 1158 g for well-watered and 854 g for drought were determined using the Decagon Soil Moisture Sensor and taking readings of fully saturated and completely dry soil. Interpolated values of 80% and 25% capacity were computed and used for the two treatments, respectively. To ensure plant viability and initial consistency, during the first four days after transplanting all plants were given water to match the well-watered weight and were also given an additional volumetric watering of 40ml once per day. On the fifth day after transplanting, drought treatments were enforced.

#### Image segmentation and feature extraction

Imaging began 2 days after planting (DAP). Every plant was imaged from two sides (0 and 90 degrees) each day, for both visible and near-infrared (NIR) cameras, totaling four images per plant per day. All images were processed using the Bellwether Workflow found in PhenotyperCV (https://github.com/jberry47/ddpsc_phenotypercv). Each image is color-corrected using a previously described algorithm (Berry et al. 2018) and has the background removed by image subtraction. To obtain a mask, a pipeline is employed that consists of a combination of: eroding, dilating, thresholding, region of interest (ROI) selection, and logical operators. Using the final mask, morphological characteristics, hue histogram and NIR histogram are extracted and written to file. The set of morphological characteristics obtained are: area, hull area, solidity, perimeter, width, height, center of mass x-coordinate (cmx), center of mass y-coordinate (cmy), number of hull vertices (hull_vertices), center of bounding ellipse x-coordinate (ex), center of bounding ellipse y-coordinate (ey), length of bounding ellipse major axis (emajor), length of bounding ellipse minor axis (eminor), angle of bounding ellipse (angle), bounding ellipse eccentricity (eccen), bounding ellipse circularity (circ), bounding ellipse roundness (round), bounding ellipse aspect ratio (ar), fractal dimension (fd), color correction strength (det), and indicator for out of frame (oof). As part of the feature extraction of the images, the NIR histogram for each image is produced. Post-processing of the histogram is done by normalizing the distribution by the size of the plant and calculating the average gray level for each image is done using weighted-mean estimation.

#### Outlier detection

Identification and removal of outliers was performed using Cook’s distance on a linear model that only included the interaction term of treatment, microbe, and time following Berry et al. (2018). This process resulted in approximately 7% of the data, 2029 images, being removed from further analysis.

#### Shapes ANOVA

To assess the variability to the drought, microbe, and interaction terms on each of the phenotypes, a fully random effect model was performed using R package lme4. For each phenotype, the sum of squares associated with each term was extracted and normalized to the total variance of the model to obtain the amount of variance explained by each component. The Pearson correlation matrix of all 20 phenotypes for all plant images on the last day was calculated and visualized using the R package corrplot. We used the plant area to estimate the effect of spatial distribution in the phenotyping growth chamber (Berry et al 2021). To aid the data exploration and visualization of raw data from PhenotyperCV and PlantCV pipelines, a shiny app (http://shiny.danforthcenter.org/PhenoAnalyzer/) was created and the plots (Fig. 2A, B, Fig. S2A, B) can be easily reproduced, along with additional analyses, using the raw data in the supplemental data files.

#### Phenotyper final harvest

On DAP23 image data was rapidly analyzed to identify outliers (outside of the 95% confidence interval based on plant area). The high-throughput phenotyping assay was concluded on DAP25 and 10 plants were randomly selected for each treatment, avoiding identified outliers. Shoot weights (fresh and dry) and root samples (root plus rhizosphere) for each selected plant were collected.

##### DNA extraction

Four 1-inch-long root sections and the attached soil were collected together for each plant. 2-ml Eppendorf Safe-Lock Tubes containing the samples were stored at −80 °C with 4 2.38-mm stainless steel beads until processing. Root and soil samples were pulverized using a TissueLyser II (Qiagen) with cold blocks cooled in liquid nitrogen (2 minutes grinding, 30 Hz, 4 times). The orientation of sample cassettes in the TissueLyser was rotated between two grindings. DNA extractions were carried out on ground root and soil samples using DNeasy PowerPlant Pro kit (Qiagen) following the manufacturer’s instruction.

#### Bacterial 16S rRNA gene sequencing

16S rDNA Pair-End (PE) amplicon sequencing on V4-V5 regions using the primers 515F (5’-GTGCCAGCMGCCGGCGGTAA-3’) and 1064R (5’-CGACRRCCATGCANCACCT-3’) was performed on the microbiome DNA samples at the University of Minnesota Genomics Center. DNA sequence data for this experiment are available at the NCBI Bioproject repository (https://dataview.ncbi.nlm.nih.gov/object/PRJNA720397?reviewer=7acfhb2i29lt6cc4eqgrspfqse). The abundance matrix, metadata and taxonomy are available at Zenodo.

#### Processing amplicon reads and designating operational taxonomic units (OTUs)

Processing of the 16S PE amplicon sequencing data was done as described in detail (Berry et al 2021). In short, the VSEARCH workflow (Rognes et al. 2016) was used to process and curate the OTU table. Quality control was run on the OTU table to remove samples and OTUs with low coverage. Samples with less than 10,000 OTUs were removed. OTUs with less than 100 or more than 200,000 reads were also removed. This yielded 92,385 OTUs. To facilitate comparisons across the OTU table and between samples, each OTU count in each sample was scaled proportionally. The taxonomic identity of each OTU is determined using both SILVA and RDP 16S databases augmented with the known 16S sequences of the individual SynCom strains. The α-diversity metric (Shannon diversity) was calculated using the diversity function from the vegan package (Oksanen et al. 2019). Spatial influence on the microbiome data was evaluated using ANOVA and spatial correction was performed using the method described in (Berry et al 2021). After spatial correction, UMAP (Uniform Manifold Approximation and Projection for Dimension Reduction) (McInnes, Healy, and Melville 2018), both unsupervised and supervised approximations, were used to assess the treatment effects on global microbiome profile. The interaction between the drought and microbial treatments was used as the supervision factor (Calibrated Abundance ~ Drought X SynCom). OTUs with significantly differentiated abundance in each microbial treatment under drought were identified using the indicspecies (indicator species) package in R (De Cáceres and Legendre 2009). The results of the indicator OTUs in different microbial treatment groups were visualized as Venn Diagram using Venny (https://bioinfogp.cnb.csic.es/tools/venny/). Calibrated abundance at the phylum level was fitted with the generalized linear mixed model based on negative binomial distribution (nb glmm) to detect the enrichment. Lsmeans function in lsmeans package in R was used to test the significance of the enrichment effects in the resulting models. P-values were adjusted using the method of Benjamini-Hochberg and FDR was controlled to the level of 0.05.

#### Cluster analysis and heat map to define the indicator OTUs

The relative abundance matrix of the indicator OTUs compared to the control (Indicator OTUs X samples) was calculated by dividing the abundance of each indicator OTU in its sample over the median abundance of that indicator OTU in the control samples. A hierarchical clustering was applied over the relative abundance matrix using the function hclust in the stats base package in R. The relative abundance matrix of the indicator OTUs was further visualized using a heatmap. The rows in the heatmap were ordered according to the dendrogram order obtained from the cluster analysis of the indicator OTUs. The heatmap was colored on the basis of the log_2_-transformed fold change compared to the control.

#### Change-point hinge models associating the plant phenotypes and microbiome abundance

Calibrated abundance of each OTU was fitted against both plant phenotypes, plant area and fresh shoot weight, with the change-point hinge model, a function provided by R chngpt package. An OTU was considered a hit if the slope of the line after the estimated threshold is significantly non-zero for both plant phenotypes. The significant OTUs were visualized using upsetplot, a function provided by R UpSetR.

### Sorghum field Experiment (Fig. 4–5, Fig. S5)

**Fig. 4.**
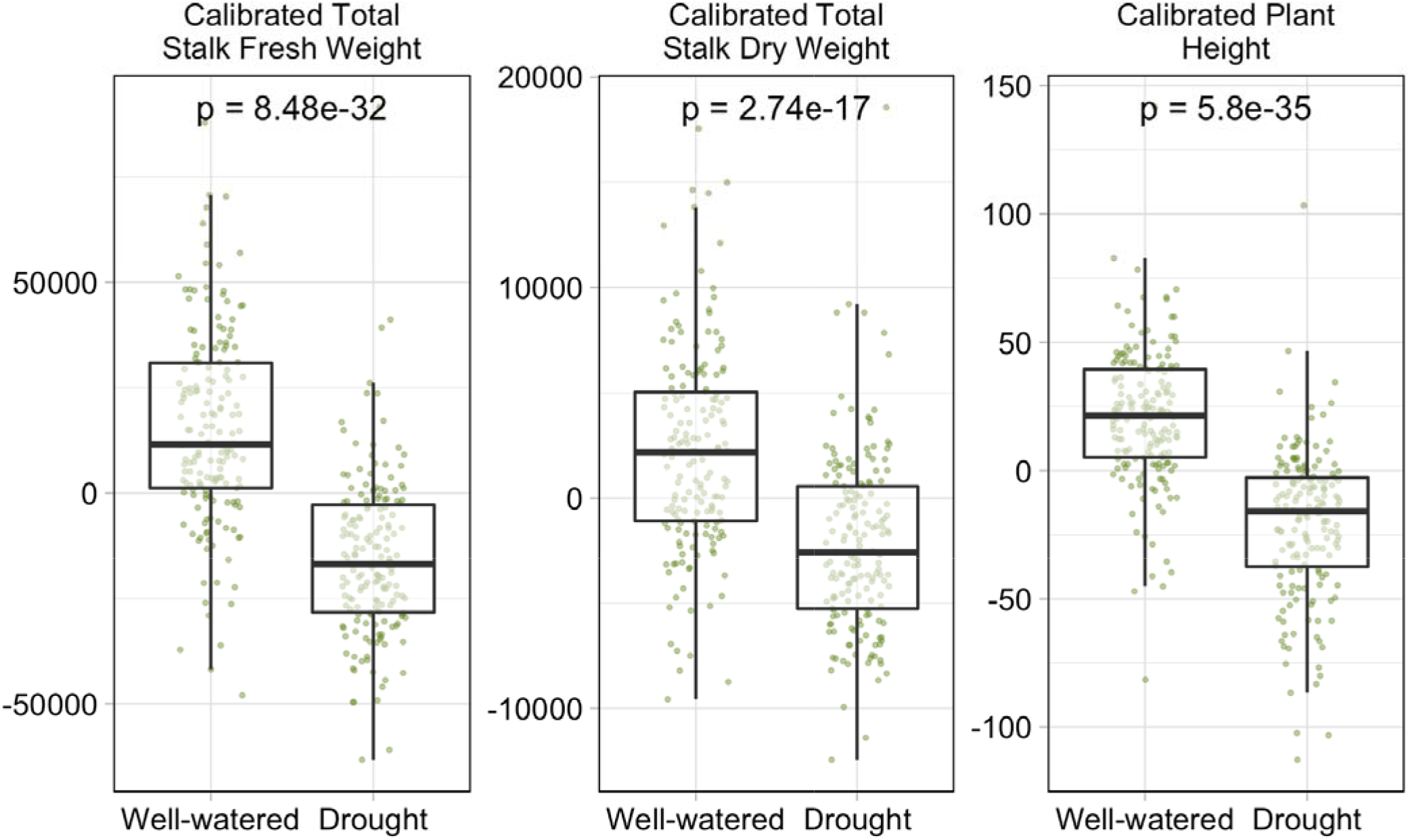
Drought treatment had negative impact on sorghum growth phenotypes in the field assay. The green dots represent the growth phenotypes of the sorghum plant samples. The horizontal bars within boxes represent medians. The tops and bottoms of the boxes represent the 75th and 25th percentiles, respectively. The upper and lower whiskers extend to data no more than 1.5× the interquartile range from the upper edge and lower edge of the box, respectively. T-tests were performed between the drought treatments with p-values showed.

**Fig. 5.**
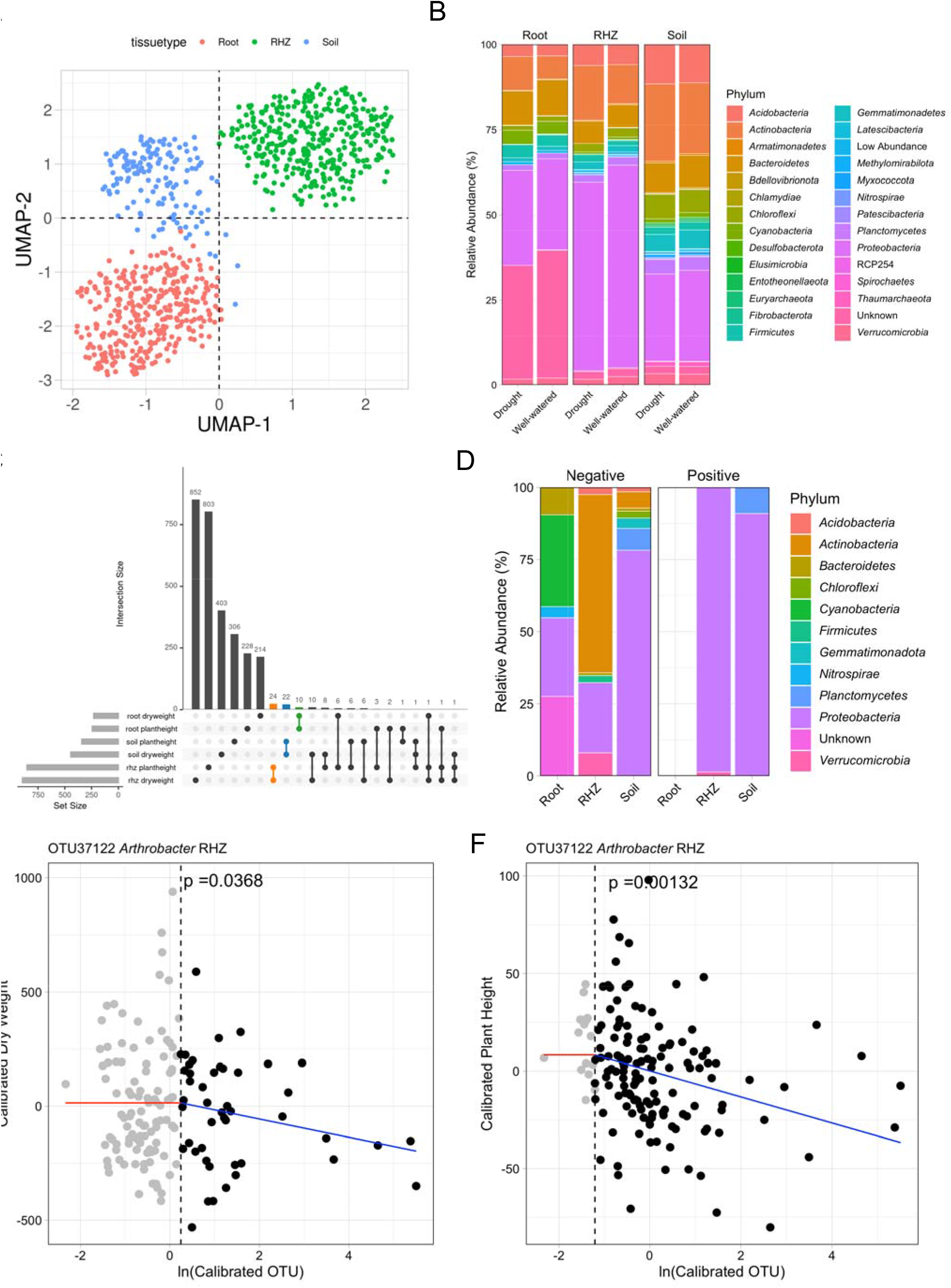
The sorghum microbiome with drought treatments in the field assay. A. The clustering of microbiome samples using unsupervised UMAP, with colors showing the tissue compartments; B. Phylum-level distribution of the sorghum microbiota with in drought treatments and tissue compartments. C. The numbers of OTUs associated to both plant phenotypes. Colors represent the OTU groups with in the same tissue compartments. D. Phylum-level distribution of the plant phenotype-associated microbiota with in the tissue compartments under drought. E, F. The change-point model fitting between OTU abundance and plant phenotypes (E, plant dry weight; F, plant height) for OTU37122 *Arthrobacter* strain. Grey dots indicate samples that did not meet the abundance threshold.

#### Field layout and experimental design

The field was located near Scottsbluff Nebraska at 41°53’39.4”N 103°41’06.1”W. The previous crop grown in this field was dry bean. Nitrogen (urea) was applied to the field area and incorporated using light tillage at a rate of 90 kg ha^−1^. Plots consisted of four rows, 76 cm apart and 4.6 m long. A split-plot design was implemented using eight replicate blocks for two watering treatments and twenty-four sorghum genotypes. The two watering treatments were delivered with a variable-rate lateral irrigation system which supplied 31.75 cm of water to the well-watered treatment and 3 cm to the water-stressed treatment. The well-watered plots were irrigated every 7-10 days and the water-stressed plots were initially irrigated to allow the crop to emerge and then irrigation was stopped. Seeds were supplied by the Kresovich, Rooney and Dweikat labs, were treated with Concep III, and were planted on June 7, 2017. Glyphosate at 1.54 kg a.i. ha^−1^ and S-metolachlorat 1.42 kg a.i. ha^−1^ were sprayed to the field one day after planting. Final biomass harvest and sampling was done on September 19, 2017.

#### Field harvest measurements and microbiome sampling

The fresh and dry weights of plots were measured by hand harvesting a 91 cm section of a row. The number of stalks was recorded, panicles were cut off and stalks and panicles, when present, were weighed separately. After weighing a 91 cm section, a subset of stalks and panicles were dried to a constant weight and a dry weight to fresh weight ratio was calculated to determine the dry weight of the entire 91 cm section. Plant heights were measured as the average height of plants in one of the two center rows of the 4-row plot with a telescoping measuring stick which could be aligned with the top of the plants. The plant phenotype data was then normalized based on a soil chemical spatial structure as described in (Berry et al 2021). The plant phenotypes were further normalized by removing the genotype effects after calibration from the soil’s chemical spatial distribution. DNA was extracted from roots, rhizosphere and the bulk soil for two plants in each plot using methods described in McPherson et al (McPherson et al. 2018) and all samples were sent for 16S rRNA gene amplicon sequencing at JGI.

#### Field sample collection and DNA extraction

The protocol previously published was used (McPherson et al. 2018). Briefly selected plants were excavated using a shovel. The excess soil (approximately 200 g) from the excavated root ball was shaken off and collected into quart-size Ziploc bags. A representative sample of root types from each plant were cut with a scissor and placed in 50 mL tubes with phosphate buffer (6.3 g L^−1^ NaH_2_PO_4_, 8.5 g L^−1^ Na_2_HPO_4_ anhydrous). After vigorous shaking, the roots were removed from the tubes and placed in new 50 mL tubes. The soil that was released from the roots (rhizosphere) was saved in the 50 mL tubes with phosphate buffer. The rhizosphere, roots and soil were placed on ice and transported to the laboratory. Solutions of sodium hypochlorite (5.25%) and ethanol (70%) were used to surface sterilize the roots for 30 sec in this respective order, followed by washing three times with sterile ultrapure water. Roots were then cut and frozen in 15 mL tubes. Liquid N was then used to grind the roots to homogenize and access the endosphere microbial communities. The rhizosphere samples were first filtered (100 μm mesh) to remove large debris, then pelleted (3000 x g for 10 min) and resuspended with 1.5 mL phosphate buffer. After transferring to a sterile 2 mL tube, the rhizosphere was re-pelleted and the supernatant was discarded. The rhizosphere pellet, the ground roots and a small sample of soil were stored in 2 mL tubes at −20 °C until DNA extraction. The remaining soil was stored in the Ziploc bags at 4 °C. The rhizosphere and bulk soil DNA extraction was performed using the MoBio PowerSoil-htp 96-well soil DNA isolation Kit, while the endosphere DNA was extracted using the Applied Biosystems (ThermoFisher Scientific) MagMax Plant DNA isolation kit. A KingFisher robot was used to automate the DNA extractions.

#### Bacterial 16S rRNA gene amplicon sequencing

DNA was quantified and then amplified in 96 well plates with single indexed primers targeting the V4 region of the bacterial 16S rRNA gene (W. Walters et al. 2016; W. A. Walters et al. 2018). Chloroplast and mitochondrial Peptide Nucleic Acid (PNA) blockers were used to prevent chloroplast and mitochondrial amplification in root endosphere samples (Lundberg et al. 2013). Amplified samples were multiplexed at 184 samples per 2 x 300bp PE Illumina MiSeq sequencing.

#### Field microbiome data analysis

Data analysis followed the same methodology described above for the Phenotyping Experiment and as described in (Berry et al 2021). This includes processing of the 16S raw reads, defining and normalizing OTUs, calculating differentially abundant OTUs and using the change point hinge models to identify positive and negative associated OTUs.

### Software, Data and code availability

Segmentation and feature extraction of the images was performed with software written in C++ that is freely available at https://github.com/jberry47/ddpsc_phenotypercv and must be compiled against OpenCV (version >= 4.0) with the extra modules: ml, aruco, and ximgproc. Additional dependencies are listed in the documentation with instructions on how to install them. Statistical analyses were performed using R version 3.5.2 (Team 2017) with the following packages: NBZIMM v1.0, lsmeans v2.30-0, emmeans v1.4.8, ggtext v0.1.0, uwot v0.1.8, forcats v0.5.0, purrr v0.3.4, readr v1.3.1, tidyr v1.1.1, tidyverse v1.3.0, data.table v1.12.8, tibble v3.0.3, doParallel v1.0.15, iterators v1.0.12, foreach v1.5.0, chngpt v2019.11-26, UpSetR v1.4.0, indicspecies v1.7.9, ggrepel v0.8.2, patchwork v1.0.1 ggsci v2.9, ggpubr v0.4.0, gdata v2.18.0, compositions v1.40-3, robustbase v0.93-5, tensorA v0.36.1, DAtest v2.7.11, vegan v2.5-6, permute v0.9-5, gridExtra v2.3, stringr v1.4.0, lme4 v1.1-23, Matrix v1.2-18, scales v1.1.1, reshape2 v1.4.4, car v3.0-9, carData v3.0-4, factoextra v1.0.7, FactoMineR v2.3, corrplot v0.84, Hmisc v4.3-0, Formula v1.2-3, survival v3.2-3, lattice v0.20-41, ggplot2 v3.3.2, plyr v1.8.6, dplyr v1.0.1, dendextend v1.13.4, ggdendro v0.1.21. Raw image data and R script(s) for all data processing and figure generations can be found at *Zenodo*.

## Results

### A synthetic community and specific *Arthrobacter* strains caused root-growth inhibition on sorghum seedlings

Previous work demonstrated that specific synthetic communities (SynComs) cause root growth inhibition (RGI) phenotypes in *Arabidopsis* (Finkel et al. 2020). To investigate whether the SynComs cause similar phenotypes in sorghum, a sorghum germination assay was performed. Three SynComs were constructed for this assay: SynCom A consisted of 24 strains from a SynCom that did not cause RGI in *Arabidopsis* (Module A in (Finkel et al. 2020)); SynCom B consisted of 29 strains selected from SynComs that did cause RGI in *Arabidopsis* (Modules C+D in (Finkel et al. 2020)); and SynCom B+V consisted of the 29 SynCom B strains plus the six *Variovorax* strains from SynCom A. Sorghum seeds were soaked in SynComs overnight and then transferred into germination pouches. The sorghum seedling roots were imaged at 4 and 7 <u>d</u>ays <u>a</u>fter <u>p</u>lanting (DAP). The results showed that compared to the controls, SynCom A- and B+V-treated sorghum seedlings had longer primary roots, while SynCom B-treated seedlings displayed the shortest roots (Fig. 1A, Fig. S1A). These data suggest that SynCom A and B promote and inhibit sorghum root growth, respectively. Considering the consortium compositions, these results also suggest that *Variovorax* strains in SynCom B+V suppress the RGI phenotype elicited by SynCom B. These results are consistent with what was previously reported for *Arabidopsis* and tomato (Finkel et al. 2020), demonstrating that this effect is consistent across both monocots and dicots.

Next, we investigated the contribution of individual strains to the RGI phenotype. Each strain was tested using the germination assay (Fig. 1B). At seven DAP, 11 strains (Fig. 1B, Table S1) out of 53 caused RGI in sorghum as compared to the control seedlings, most of which belonged to SynCom B. In addition, this assay revealed that eight SynCom A strains can promote root growth, including two *Variovorax* strains. Out of the 53 strains tested, 14 were previously reported to cause RGI in *Arabidopsis*. Among those, only two strains can also induce short roots in sorghum, both of which were *Arthrobacter* strains (Fig. 1B, Table S1).

Despite statistically significant effects on root growth, we also observed a large amount of variation among experimental replicates. We hypothesized that this may be due to the level and/or location of colonization. Thus, to further investigate bacterial colonization on the sorghum roots we selected two representative strains: *Variovorax* strain CL14, the root growth promoter (the leftmost strain in Fig. 1B), and *Arthrobacter* strain CL28, the root growth suppressor (the third rightmost strain in Fig. 1B) (Fig. S1B). These strains were applied to sorghum seeds and root length was measured at 7 days. In addition, bacterial populations on the root surface and within the root tip were quantified. The results suggest that the *Variovorax* strain CL14 is a robust rhizosphere colonizer (Fig. S1B right). Endosphere colonization of CL14 was only observed in a few of the sorghum roots (green squares) and notably, these replicates showed shorter root lengths compared to rhizosphere colonized roots (green triangles). The control group also had replicates with short roots, suggesting that this short root phenotype may be independent of microbial treatment. *Arthobacter* strain CL28 also colonized the rhizosphere for the majority of replicates, with only a few replicates showing endosphere colonization (Fig. S1B left). Among the replicates with rhizosphere CL28 colonization, we observed a trend towards shorter sorghum root length. This suggests that there may be a dose-dependent effect for the strength of RGI by CL28. Taken together, these data support our hypothesis that the variation observed in Figure 1 may be at least partially explained by the location and level of colonization, even in this relatively controlled experimental system.

### Drought and microbial treatments alter sorghum growth in a high-throughput, controlled environment experiment

Based on the data in Figure 1A, we hypothesized that SynCom B-treated sorghum would show increased susceptibility to abiotic stresses such as drought and that SynCom A-treated sorghum plants would show relative tolerance because of their root length phenotypes. In addition, we wanted to investigate whether the SynCom-mediated phenotypes would transfer to more complex non-sterile environmental conditions. To address these questions, we performed a 24-day-long experiment using the high-throughput Bellwether phenotyping platform (Lemnatec system) and measured the aboveground phenotypic effects on sorghum growth of the SynComs across the well-watered and drought conditions.

Sorghum seeds were germinated in the presence of microbes, planted in steam sterilized soil and loaded onto the phenotyping platform. All pots were well-watered for four days prior to starting the drought treatment. Every plant was weighed and then watered if necessary (below target weight) and imaged each day for a total of 24 days. Both RGB and near-infrared (NIR) images were collected. NIR intensity may be used as a proxy for water content wherein a lower value correlates with higher plant water content (Fahlgren, Gehan, and Baxter 2015). At the end of the experiment, fresh and dry shoot weights were quantified. In our experiment, 80% of the variance in plant area was explained by the treatment factors, meaning plant area robustly responded to the treatments (Fig. S2A). Plotting plant size and NIR intensity over time revealed several striking differences (Fig. 2A, B). First, a clear drought treatment effect was observed for the control (no microbial seed treatment) plants as measured by reductions in plant area and increased NIR intensity, and this correlated with fresh and dry shoot biomass at the end of the experiment (Fig. 2C, Fig. S2C). This strong correlation between plant area and biomass is consistent with previous reports (Veley et al. 2017; Berry et al. 2018). In addition, considering the microbe treatments, we observed that under drought conditions, SynCom A- and SynCom B+V-treated plants performed better than control plants. Further, SynCom B-treated plants performed worse under drought than control plants. These patterns were observed for plant area, NIR intensity and shoot fresh weight at the end of the experiment (Fig. 2). We also considered shoot dry weight at the end of the experiment and while similar trends were observed, the differences were not significant (Fig. S2C).

### *Arthrobacter* and *Variovorax* strains applied to sorghum seeds colonize and persist in sorghum roots

The observed phenotypic differences among the microbial treatments suggests that the microbes in SynCom A and SynCom B+V protect sorghum from drought stress while the microbes in SynCom B exacerbate the negative impacts of drought. To test whether the microbes applied at the beginning of the experiment were able to persist with the sorghum plants and to potentially pinpoint specific microbes within each SynCom with major roles in affecting above ground phenotypes, we characterized the root microbiomes of each plant using 16S rRNA gene amplicon sequencing.

The vsearch workflow was used to cluster 16S rRNA gene amplicon sequences at 99.5% identity into operational taxonomic units (OTUs). A total of 7904 distinct OTUs were observed after quality filtering. Of the 53 individual strains that make up the three SynComs, only five strains had corresponding OTUs that were detected at the end of the experiment. This result demonstrates that not all the SynCom strains persisted through the phenotyping assay. This also showed that the majority of OTUs present at the end of the phenotyping assay originated from the non-sterile environment in which the experiment was performed.

Although the phenotyping experiment was performed within a controlled environment growth chamber, spatial effects may still occur and can introduce unwanted experimental noise. Indeed, although treatments were randomly distributed throughout all four greenhouses, greenhouse 1 (GH1) appeared to have smaller plants than the other greenhouses. Therefore, both plant area and the OTU table were adjusted for spatial effects as described (Materials and Methods and Berry et al 2021). Comparison of the original and calibrated data showed that while plant area did not change significantly, the OTU table did show significant differences (ANOVA, p-value = 2.2 × 10^−16^) and therefore, the calibrated table was used for subsequent analyses (Fig. S3A). To assess the effect of drought conditions and microbial treatments on the global microbiome profiles, we considered an unsupervised and a supervised uniform manifold projection (UMAP) analysis. While a strong drought treatment effect on the microbiome was observed with both approaches (Fig. 3A, Fig. S3B), only the supervised UMAP was able to detect a microbe treatment effect on the microbiomes, which is consistent with a strong effect from the environmental (non SynCom-derived) microbiome. To test whether drought influenced the diversity of microbial communities, we considered the Shannon diversity among the different treatment groups. SynCom A- and B+V-treated samples had significantly lower Shannon diversity compared to that of the SynCom B-treated samples, suggesting the SynCom A and B+V treatments decreased the richness and evenness of the sorghum rhizosphere microbiome (Fig. S3C). Comparing the microbiomes at the phylum level revealed several groups of microbes that were differentially abundant between the treatments (Fig. 3B). For example, *Actinobacteria* (which includes *Arthrobacter*) were more abundant in the drought treated samples (Fig. S3D).

To identify specific OTUs that were enriched among the microbe treatment groups under drought, we employed the ‘indicator species’ algorithm (De Cáceres and Legendre 2009). The resulting lists of OTUs were compared to the SynCom starting inoculums to look for overlap (Fig. S4). For example, SynCom A inoculum comprises 18 unique strains (Fig. S4A, upper Venn diagram). At the end of the experiment, we observed 5 OTUs that were specific to SynCom A-treated plants (Fig. S4A, lower Venn diagram) and indeed, the taxonomic annotations of these 5 OTUs were among the original list of SynCom A strains. In this manner, across all treatments, 18 OTUs were identified as likely SynCom-derived OTUs and in each case, the OTUs were either unique to, or strongly enriched in the samples in which they were inoculated (Fig. S4B). These OTUs represented 1%, 2.2% and 2.6% of the relative abundance in SynCom A-, B- and B+V-treated plants, respectively. These data indicate that only a subset of the SynCom strains was able to persist throughout the experiment and again highlight the significant environmental component of the microbiome.

Based on recent reports on *Arthrobacter* and *Variovorax* and our sorghum seedling data (Fig. 1), we were particularly interested in OTUs corresponding to these two genera. Under drought, SynCom A and SynCom B+V shared one enriched OTU which corresponded to *Variovorax* (OTU148636) (Fig. 3C, Fig. S4B). SynCom B and SynCom B+V shared 13 enriched OTUs, all of which had matches from the starting inoculums and one of which corresponded to *Arthrobacter* (OTU194097) (Fig. 3C, Fig. S4B). We note that this OTU was also detected in SynCom A/drought treated plants, albeit at a lower abundance possibly suggesting some level of contamination between the treatments or a similar microbe present in the environment. These data suggest that *Arthrobacter* and *Variovorax*, applied to sorghum seeds, were able to persist with developing sorghum roots over the course of the four-week experiment.

Further, these analyses revealed many additional OTUs, presumably from the non-sterile environment in which these experiments were performed, that were specifically enriched or depleted in the presence of specific SynComs. Most strikingly, a large group of OTUs was depleted during drought stress and in the presence of SynCom A (Fig. S4B). These results suggest that not only can many of the SynCom strains persist in a complex environment, they may also dramatically shape the resulting microbiome in a stress responsive manner.

### Colonization by *Arthrobacter* and *Variovorax* strains correlate with increased and reduced sensitivity to drought, respectively

Next, we queried the dataset for OTUs whose abundance correlated with plant phenotypes under drought, regardless of the microbial treatments. We reasoned that a given microbe may only influence plant phenotype once a certain amount of colonization was achieved. Change-point models accommodate this concept by allowing for no effect on the plant phenotype until a certain abundance threshold is reached, after which a linear trend between quantity of a microbe and phenotype is observed. A microbe is considered a “hit” for having a significant impact on a plant phenotype, if the regression slope after the estimated threshold is significantly non-zero, either negative or positive. Further, to reduce the amount of false-positive hits we assessed two plant phenotypes, plant area and fresh shoot weight, for every microbe. To qualify as a ‘hit’, the OTU had to exhibit significance in both phenotypes in the same direction.

In total, 209 and 89 OTUs, within the whole OTU table, were negatively and positively associated with both plant phenotypes under drought, respectively (Fig. 3D, Table S2 and S3). The relative abundance of plant phenotype associated OTUs at the phylum level were distinct between positive and negative associations (Fig. 3E). The OTUs that were positively associated with plant growth were more likely to be *Bacteroidetes* (adjusted p value = 2.16*10^−5^) and less likely to be *Firmicutes* (adjusted p value = 2.1*10^−33^) than those negatively associated with plant growth. These results suggest that bacteria within the *Bacteroidetes* and *Firmicutes* phyla may have positive and negative effects on plant growth, respectively.

We cross-referenced the plant phenotype associated OTUs (Fig. 3D) with those that showed differential abundance during drought treatment among the four microbial treatments (Fig. S4B). This yielded eight OTUs, all of which negatively affected plant phenotype and seven of which (all non-SynCom-derived) were depleted in the SynCom A samples under drought. This suggests that SynCom A treatment may decrease the abundance of deleterious environmental strains under drought conditions. Further, the eighth OTU identified from the two lists, OTU194097 (*Arthrobacter*), was among the inoculum strains for SynCom B and B+V and showed a significant negative correlation with plant phenotypes (p-value = 0.003, *R^2^* = 0.907 for plant area; p-value = 0.017, *R^2^* = 0.792 for fresh shoot weight) (Fig. 3F and G). Combining these results with those from the sorghum seedling assay, we conclude that *Arthrobacter* strains are deleterious for sorghum growth under drought stress. Notably, the five *Arthrobacter* strains evaluated in Fig. 1 all cluster into a single OTU - OTU194097.

### *Arthrobacter* strains negatively impact sorghum under drought conditions in the field

In parallel to the high-throughput phenotyping experiment described above, we performed a large-scale field experiment. In 2017, twenty-four varieties of sorghum were evaluated for performance across well-watered and drought conditions. Multiple sets of data were collected, including plant traits at the final harvest (plant height, fresh and dry stalk weight, and panicle weight), soil chemical content and properties (calcium, magnesium, phosphate levels and etc.), and microbiome samplings from three belowground compartments for each plot (root, rhizosphere and soil). Initial analyses revealed strong evidence of heterogeneous spatial distribution of soil factors. Therefore, all plant phenotypes were calibrated to exclude soil nutrient and spatial effects. This approach is described in detail in Berry et al (Method paper citation). In brief, soil nutrients were dimension reduced to the first three principal components and were regressed against using a linear model that included the spatial covariance structure. Calibrated phenotypes were the raw residuals of this model and were used in the subsequent analysis having accounted for these covariates. Further, all the plant phenotypes were adjusted for genotype effects using linear regression and retaining the residuals. After soil factor calibration and genotype adjustment, plant biomass phenotypes, including plant stalk weight and plant height, all suggested that the drought treatments had impaired the plant growth (Fig. 4), which validated the drought treatments.

To investigate the microbiome composition associated with each plant, DNA was extracted from three compartment samples (root, rhizosphere (rhz), and soil) and analyzed. After quality control, the OTU table was calibrated to account for the spatial soil property effects (Berry et al 2021). To visualize general relatedness between the sample types, we considered an unsupervised UMAP analysis. The data clustered most strongly by tissue compartment (Fig. 5A). We also considered a supervised UMAP and observed that the drought treatment effect was most obvious in the rhizosphere samples, consistent with previous reports (Fig. S5)(Naylor et al. 2017; Xu et al. 2018). Also consistent with previous reports, the Shannon diversity of the microbiome was lowest in the root samples. We did not observe a drought impact on Shannon diversity possibly because all genotypes are collapsed in this analysis (Fig. S5C). *Proteobacteria*, *Actinobacteria*, *Bacteroidetes* and *Acidobacteria* were the most abundant phyla in all samples (Fig. 5B). Comparing the relative abundance in different compartments, both *Actinobacteria* and *Acidobacteria* were highest in soil and lowest in root, while *Proteobacteria* was highest in rhz and lowest in soil, and *Bacteroidetes* was highest in root and lowest in rhz. Moreover, *Chloroflexi* was much higher in soil than in rhz and root. Compared to those in well-watered conditions, the relative abundance of *Actinobacteria* increased in drought conditions, while other major phyla remained similar.

We used a zero-inflated negative-binomial generalized linear mixed model (ZINBGLMM) to identify significantly enriched and depleted OTUs between the well-watered and drought conditions for each sample type (FDR was controlled to 0.05) (Fig S5D). The number of differentially abundant OTUs, as a proportion of the total number of OTUs for each compartment, was smallest in the soil samples (Fig. S5D). These observations suggest that the plant roots actively modulate the root and rhizosphere microbiome in response to abiotic stresses such as drought, consistent with previous reports (Naylor et al. 2017; Fitzpatrick et al. 2018; Xu et al. 2018). *Actinobacteria* were enriched under drought in all three compartments (Fig. S5E), and the genus *Arthrobacter* was among the enriched Actinobacterial genera (Fig. S5F).

As a final step, we applied a similar methodology used for the high-throughput phenotyping study described above to identify putative phenotype-associated microbes from the field data. We queried the field data using change-point hinge modeling with two plant phenotypes, plant height and stalk dry weight. Once soil property effects were removed from the OTU table and plant phenotypes, the analysis revealed 22, 24 and 10 phenotype associated OTUs from soil, rhizosphere and root, respectively (Fig. 5C) and these OTUs were further broken down into positive and negative associations with both phenotypes (48 OTUs in total) (Fig. 5D). OTU37122 corresponding to *Arthrobacter*, was among the negative plant-phenotype-associated OTUs identified from the rhizosphere-derived samples (Fig. 5E and F). This OTU was also enriched under drought conditions (Fig. S5E and G). While *Variovorax* was not among the OTUs that positively correlate with plant phenotype, this analysis did reveal several other candidate beneficial bacteria. Cross referencing this list with positively associated OTUs from the high-throughput phenotyper experiment revealed two closely related OTUs. Based on the NCBI 16S rRNA database, these OTUs most closely match *Nordella oligomobilis* (*La Scola, Barrassi, and Raoult 2004*), within the *Rhizobiales* Order (Fig. S5H, Table S4).

## Discussion

Drought is one of the most serious and unpredictable challenges associated with modern day farming. Exacerbated by the effects of climate change, farmers without easy access to irrigation increasingly experience crop loss from lack of rainfall (Jaleel et al. 2009; Assefa, Staggenborg, and Prasad 2010; Daryanto, Wang, and Jacinthe 2017). Beneficial microbes are often touted as a potential method of providing crops with enhanced drought tolerance (Kim et al. 2012; Kumar and Verma 2018; Kour et al. 2019, 255; Ulrich et al. 2019). However, while many candidate beneficial microbes show promise within controlled settings, researchers struggle to translate these candidates to the field (Fukami et al. 2016). Similarly, while native soils are rich in microbial diversity, it has proved challenging to isolate individual bacteria or consortia that are beneficial when reapplied in a field setting (Finkel et al. 2017; Afzal et al. 2019). Here, we describe a three-pronged experimental approach to identifying microbes that affect sorghum drought stress tolerance.

First, we tested synthetic communities (SynComs) of bacteria designed based on interactions with the model plant, *Arabidopsis* (Finkel et al. 2020). When applied to sorghum roots, the synthetic communities elicited phenotypes very similar to those observed from *Arabidopsis* (Fig. 1A). However, individual strains were less consistent in their effects. That is, some strains that caused a short root phenotype on *Arabidopsis* did not affect sorghum and vice versa (Fig. 1B). These results may indicate some amount of host specificity (Chai et al. 2021) or simply reflect differences in the experimental assays used for *Arabidopsis* and sorghum. Regardless, these assays pointed to *Arthrobacter* and *Variovorax* as being particularly impactful on sorghum roots, similar to what was recently reported for *Arabidopsis* (Finkel et al. 2020).

Next, we tested the SynComs on sorghum over the course of a 24-day high-throughput phenotyping assay. This system was relatively uncontrolled, compared to the seedling assays, and yet robust SynCom dependent phenotypes were observed (Fig. 2). Specifically, seeds treated with the SynCom that contained *Arthrobacter* but not *Variovorax*, showed increased sensitivity to drought stress. To further understand this observation, we performed 16S rRNA amplicon sequencing and confirmed that both *Arthrobacter* and *Variovorax* had persisted with sorghum throughout the course of the experiment at least in some replicates (Fig. 3C). In searching for root-associated microbes whose abundance correlates with above ground phenotypes, we reasoned that colonization and persistence within this experimental system was likely to be non-uniform. Similarly, we reasoned that microbes may only cause impact above a certain level of abundance. To accommodate these potential sources of noise, we applied a change-point model that looks for an effect only once a certain abundance threshold is met. From this analysis, we observed strong negative correlations between *Arthrobacter* abundance and two distinct measures of plant size (Fig. 3F and G). Based on the results from the seedling assay (Fig. 1) we hypothesize that the observed phenotypes were due to a root developmental defect caused by *Arthrobacter* and/or other deleterious strains. In the future, we look forward to “in soil” advanced root imaging capability that will provide additional insight into root development during interaction with various microbes and abiotic stresses (Jiang et al. 2019).

In parallel, we undertook a large field experiment wherein drought stress was applied to various sorghum genotypes. We reasoned that because sorghum is a drought-tolerant plant, we would be most likely to isolate microbes that promote drought tolerance from drought treated sorghum, as has been broadly proposed (Santos-Medellín et al. 2017; Timm et al. 2018). Our initial attempts at analyzing field phenotyping and 16S rRNA gene sequencing data from the field revealed an abundance of experimental noise, typical of field experiments. Upon closer examination, we observed strong spatial effects for multiple measured soil chemical properties and so developed a method of accounting for this variability within our models (Berry et al 2021). We next applied the change-point model to look for OTUs that show positive or negative associations with the measured plant phenotypes (Fig. S5H). *Arthrobacter* was among the identified negatively associated OTUs in the rhizosphere (Fig. 5E and F), suggesting that this genus is likely a significant impediment to plant productivity in the field, especially under drought stress. This piece of data is also consistent with our findings that the rhizosphere colonizer *Arthrobacter* strain CL28 from *Arabidopsis* SynCom (SynCom B) inhibited sorghum root growth (Fig. S1B). While we did not observe significant correlations for OTUs that correspond to *Variovorax* in any analyses for this field, we did identify 10 OTUs that showed positive correlation with plant phenotypes including 3 OTUs that fall in the Order *Rhizobiales*, which includes previously described beneficial and pathogenic microbes (Carvalho et al. 2010).

When cross referencing the plant growth associated OTU lists from the phenotyping and field assays, we noticed a common ‘hit,’ most similar to the previously described *Nordella oligomobilis*, that is positively correlated with plant phenotypes. Little is known about this type of bacteria; however, it falls within the Order *Rhizobiales* (La Scola, Barrassi, and Raoult 2004) along with multiple OTUs corresponding to genera within Family *Bradyrhizobiaceae*. This family of *Alphaproteobacteria*, especially the Genus *Bradyrhizobium*, includes many slow-growing symbiotic rhizobial strains, many of which are beneficial for their host plants by forming nitrogen-fixing nodules. Recent studies suggested that many non-symbiotic *Bradyrhizobium* species are ecologically important for the soil microbiota and such ecotypes dominate the coniferous forest soil (VanInsberghe et al. 2015; Jones et al. 2016; Ormeño-Orrillo and Martínez-Romero 2019). Moreover, *Bradyrhizobium* strains were reported previously to degrade auxin (Egebo, Nielsen, and Jochimsen 1991; Jensen et al. 1995; Torres et al. 2021), whose level plays an important role in plant resilience (Finkel et al. 2020). Thus, it is tempting to speculate that these newly observed beneficial *Bradyrhizobiaceae* strains may have plant growth promoting properties including auxin degradation, which complement the absence of *Variovorax* in this soil.

Six common genera were negatively associated with plant phenotypes under drought in both assays (field and phenotyping assay), including *Arthrobacter, Marmoricola, Noviherbaspirillum*, *Paenibacillus*, *Pseudolabrys* and *Pseudomonas*. *Arthrobacter*, *Marmoricola* and *Paenibacillus* are gram-positive genera but have not been extensively characterized (Grady et al. 2016; Tchuisseu Tchakounté et al. 2018). Consistent with previous studies (Naylor et al. 2017; Fitzpatrick et al. 2018; Xu et al. 2018), we observed an enrichment of *Actinobacteria* under drought stress (Fig. S3D and S5E) and further presented evidence that *Arthrobacter* strains suppressed plant growth, especially under drought stress (Fig. 3F, 3G, 5E, 5F). It is well known that *Pseudomonas* strains have diverse effects on plant growth, including plant growth promotion from *Pseudomonas putida* and *Pseudomonas fluorescens* strains and plant diseases caused by pathogenic *Pseudomonas syringae* strains (Ganeshan and Manoj Kumar 2005; Trivedi, Spann, and Wang 2011; Bernal et al. 2017; Xin, Kvitko, and He 2018). In general, our results suggest that under drought stress, *Pseudomonas* is detrimental to sorghum growth. Other than *Pseudomonas*, the remaining genera are relatively understudied (Kämpfer et al. 2006; Ishii et al. 2017) and so future work will focus on culturing these bacteria and then investigating these strains for potential mechanisms of plant growth suppression. Having cyro-preserved a portion of each field derived root sample, our next endeavor will be to isolate and test these specific candidates.

In conclusion, we have demonstrated that specific isolates of *Arthrobacter and Variovorax* that affect dicot root growth also affect root growth in sorghum, a monocot (Fig. 6). Through a three-pronged approach that spanned sterile, controlled environment and field experiments, we have identified a high confidence list of novel candidate beneficial microbes. This systems-level approach allowed us to mitigate significant environmental noise to reveal underlying robust biological interactions.

**Fig. 6.**
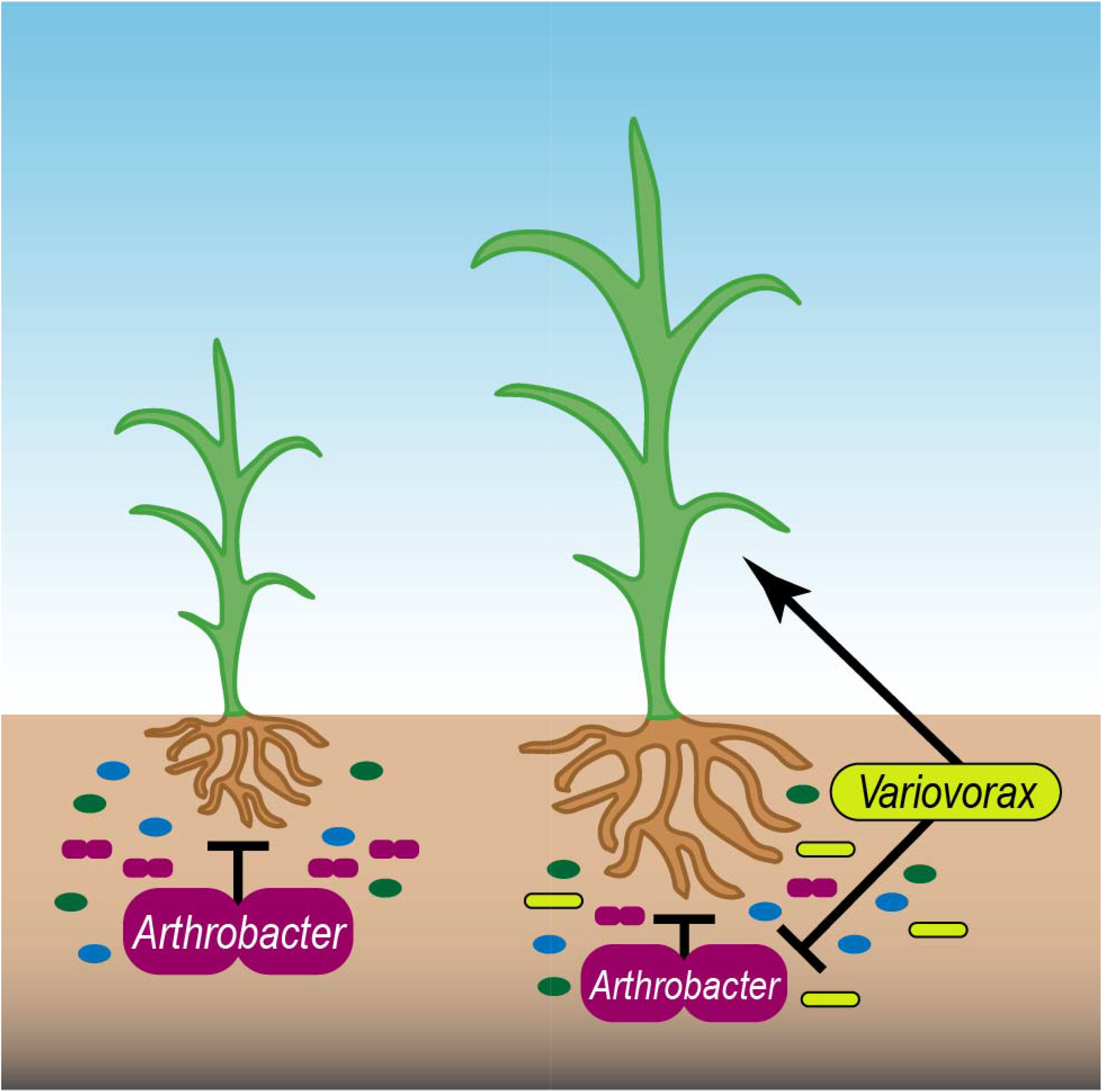
The working model of sorghum-microbiome interactions. *Arthrobacter* strains in the microbiome associated to sorghum roots have negative impact on sorghum growth, especially under stresses (left panel). *Variovorax* strains protect the sorghum growth through suppressing the activity of *Arthrobacter* strains.

## Acknowledgement

This work was primarily funded by the Department of Energy (DE-SC0014395). Additional support came from NSF-REU (DBI-1156581); NSF grant IOS-1917270 to JLD. JLD is an Investigator of the Howard Hughes Medical Institute, supported by the HHMI. OMF was supported by NIH NRSA Fellowship F32-GM117758; PL is supported by the Iowa State University Plant Sciences Institute Scholars Program; 16S rRNA gene sequencing of field samples was supported by a Community Science Program award to DPS and SGT from the DOE Joint Genome Institute. The work conducted by the U.S. Department of Energy Joint Genome Institute, a DOE Office of Science User Facility, is supported by the Office of Science of the U.S. Department of Energy under Contract No. DE-AC02-05CH11231.

---

**Fig. S1.**
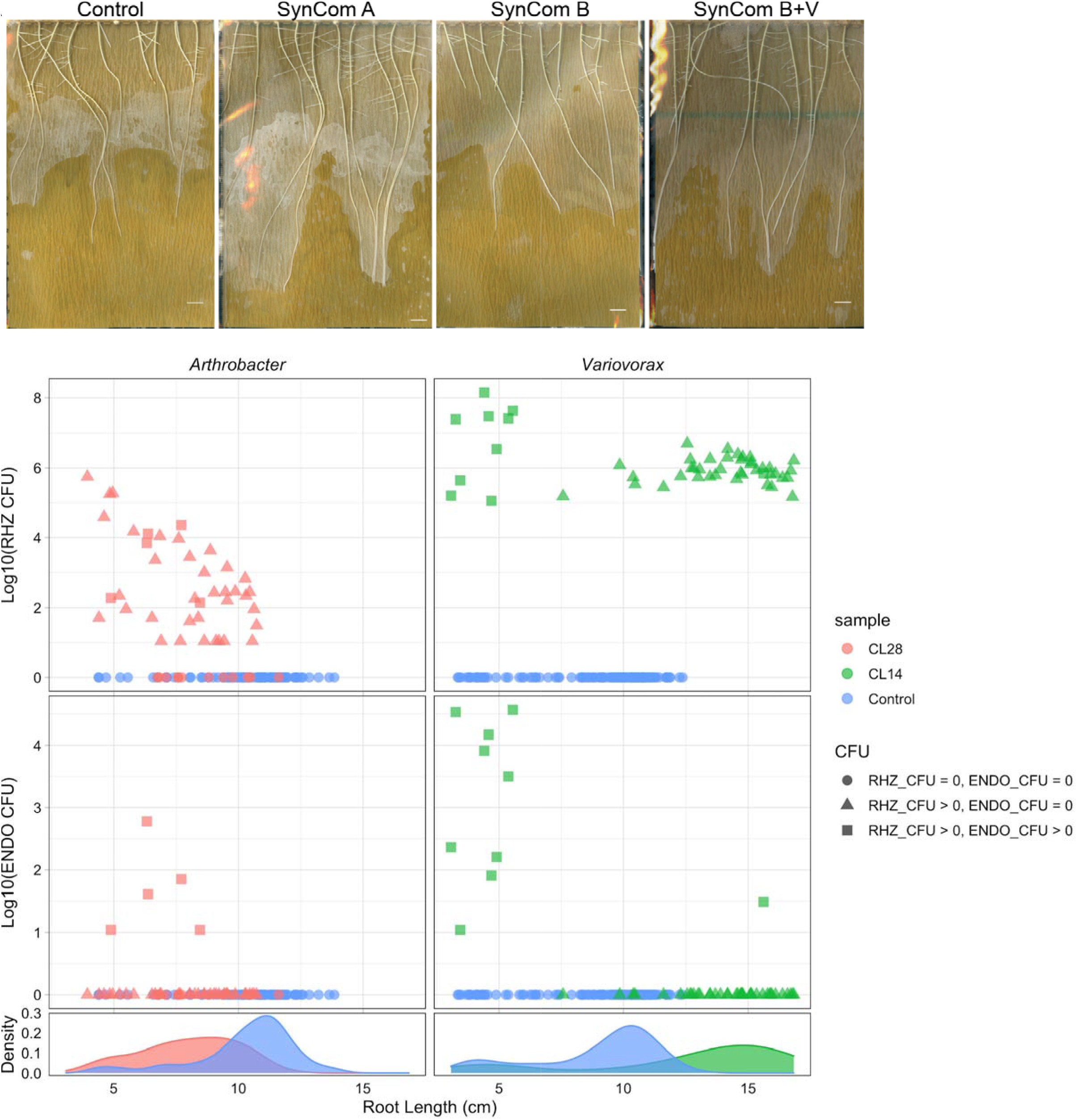
Sorghum root growth in germination pouches at DAP7 with SynCom (A) and individual strain (B) inoculations. A. Pouch replicate n=3. Bar: 1 cm. B. Colors and shapes represented the bacterial inoculations and colonization tissue compartments, respectively. RHZ = rhizosphere colonization; ENDO = endosphere colonization. CL28: *Arthrobacter* strain; CL14: *Variovorax* strain. Control = no microbial treatment. Strain details can be found in Table S1.

**Fig. S2.**
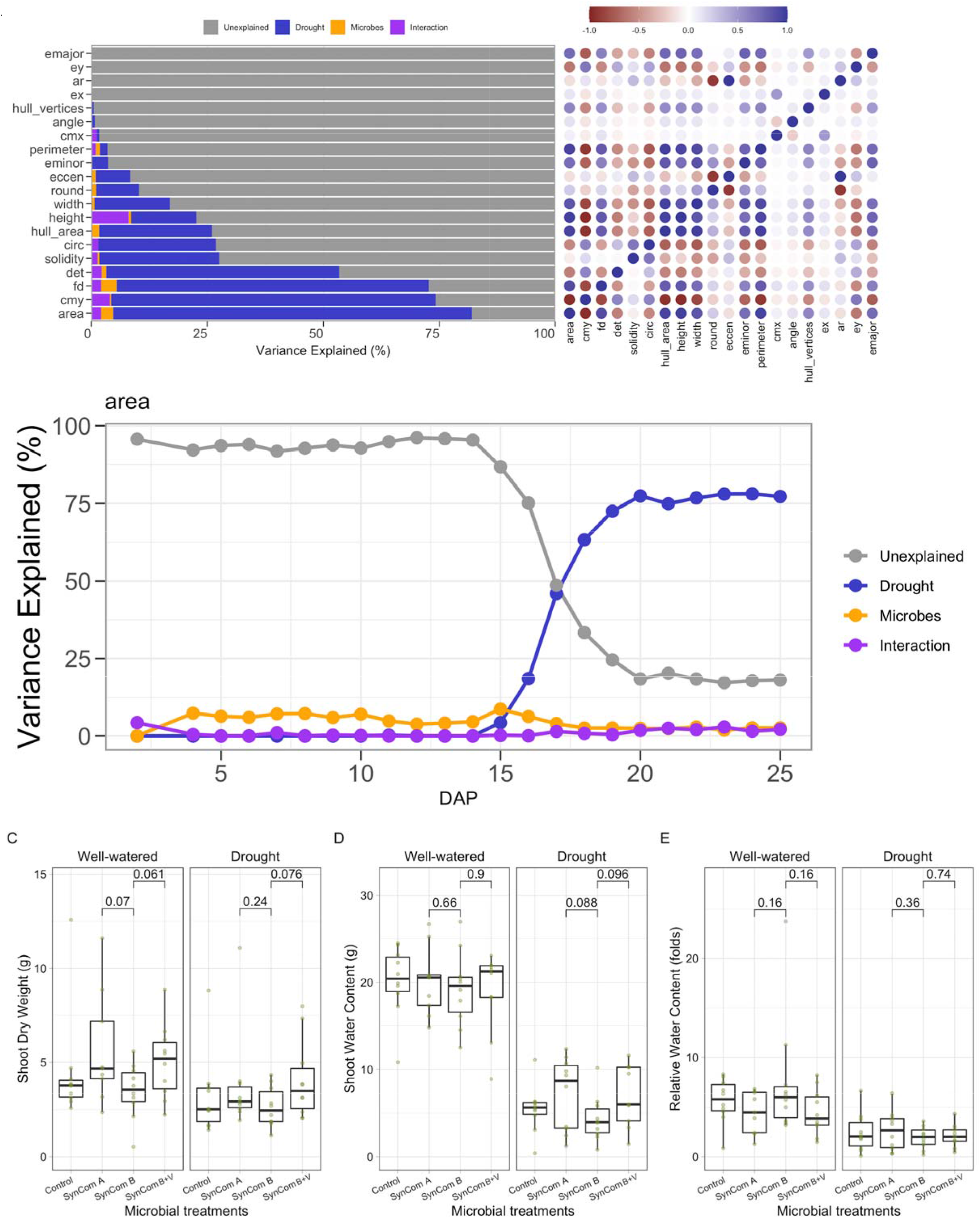
Sorghum growth phenotypes in the high-throughput phenotyping assay. A. PlantCV image analyses pipeline reports variances of 20 plant phenotypes determined with ANOVA (left), and the correlation matrix among them (right). B. The temporal changes of the variance sources in plant area. C-E. The green dots represent the shoot dry weight (C), shoot water content (D) and plant relative water content (E) of sorghum at the conclusion of the assay, respectively. Box plots display medians (horizontal line) the 75th and 25th percentiles (top and bottom of box) and the upper and lower whiskers extend to data no more than 1.5× the interquartile range from the upper edge and lower edge of the box, respectively. Pairwise t-tests were performed between microbial treatments for well-watered and drought conditions. P-values for select comparisons are shown and all others were not significant at the alpha of 0.05. The number of replicated samples for each treatment n = 10 (C-E).

**Fig. S3.**
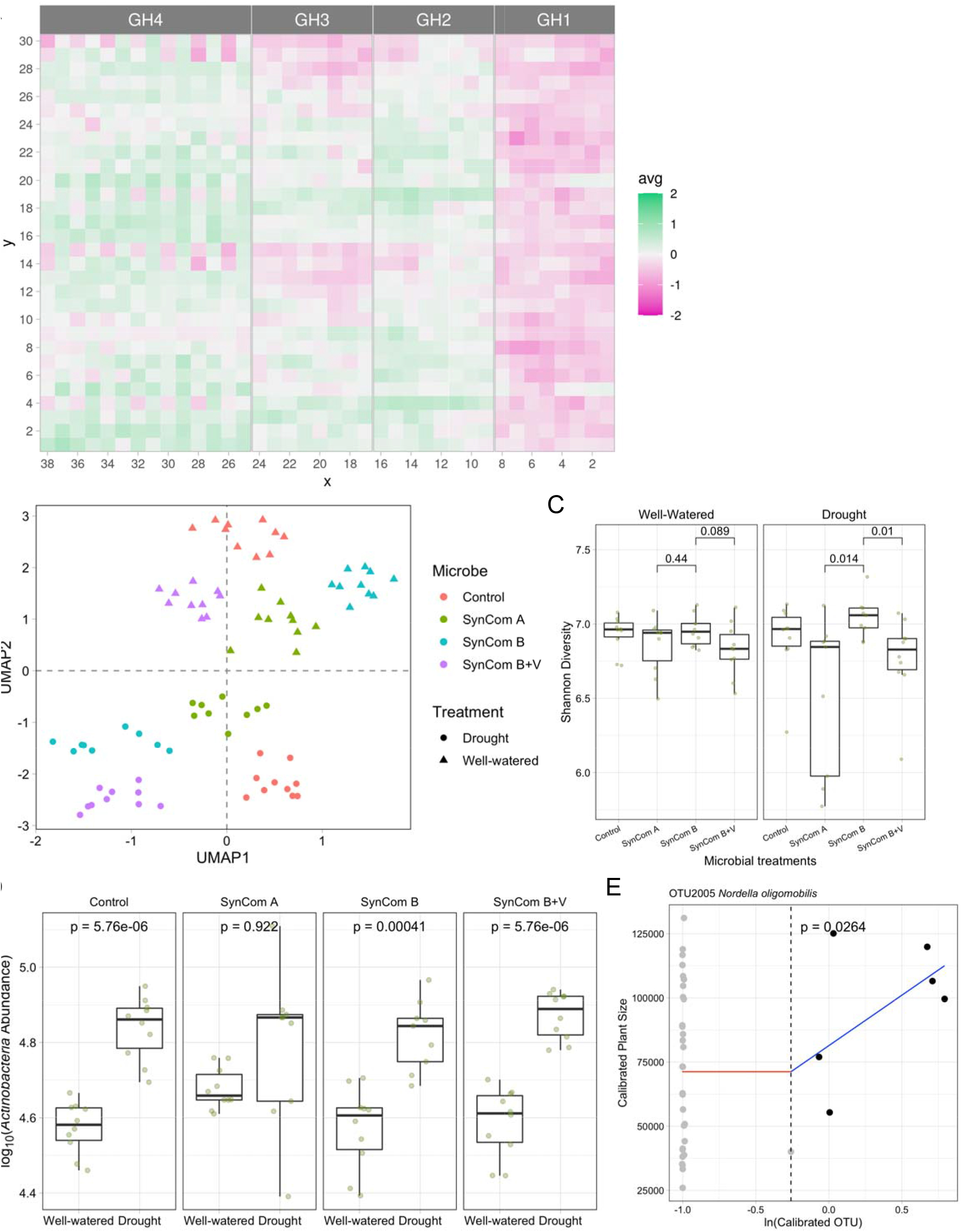
Sorghum root-associated microbiome with SynCom and drought treatments in the high-throughput phenotyping assay. A. The spatial distribution of sorghum plant size in the phenotyping facility. GH: greenhouse; avg: Average plant area. B. The clustering of microbiome samples using supervised UMAP, with colors and shapes showing the drought and microbial treatments, respectively. C. The green dots represent the Shannon diversity of sorghum microbiome samples. Pairwise Wilcoxon tests were performed between microbial treatments for well-watered and drought conditions. P-values for select comparisons are shown and all others were not significant at the alpha of 0.05. D. The abundance of *Actinobacteria* strains was enriched under drought in three of four microbial treatments. The colors represent the drought treatments. NBGLMM models were fitted between the drought treatments with corrected p-values shown. E. Example of OTU abundance positively correlating with plant phenotype based on the change point hinge model. C and D, The horizontal bars within boxes represent medians. The tops and bottoms of the boxes represent the 75th and 25th percentiles, respectively. The upper and lower whiskers extend to data no more than 1.5× the interquartile range from the upper edge and lower edge of the box, respectively. The number of replicated samples for each treatment n = 10.

**Fig. S4.**
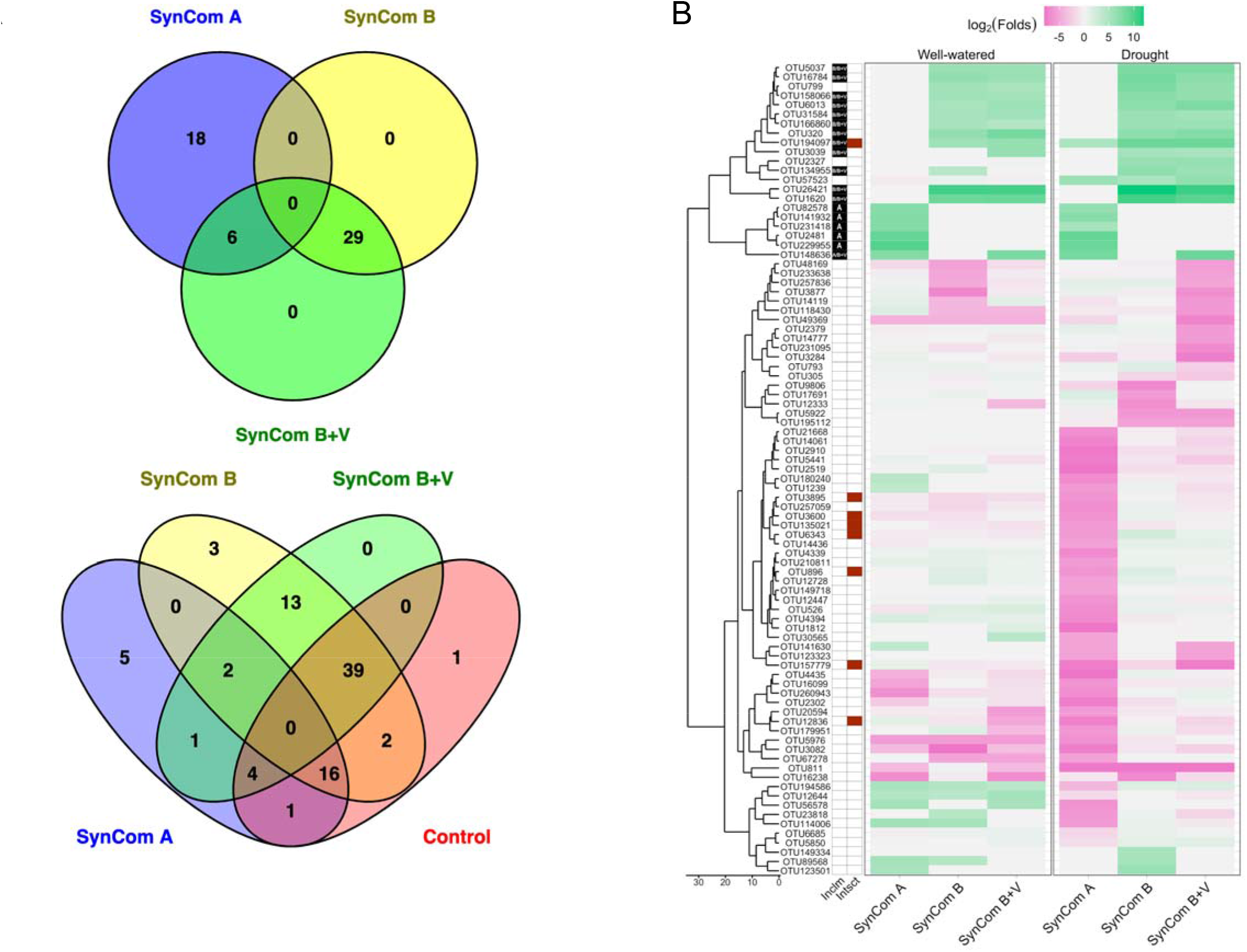
The specific OTUs enriched among the microbial treatments under drought. A. The strains shared in the inoculums (upper) and the OTUs shared in the microbiome samples (lower). B. The relative abundance changes of the significant OTUs in the microbiome samples with SynCom inoculations compared to the control. The OTU clusters in the dendrogram (left) were determined according to the OTU abundance profile across the samples. Inclm (black with SynCom signs): SynCom-derived OTUs; Intsct (dark red): OTUs with specific abundance enrichments and negatively associated to the plant growth phenotypes.

**Fig. S5.**
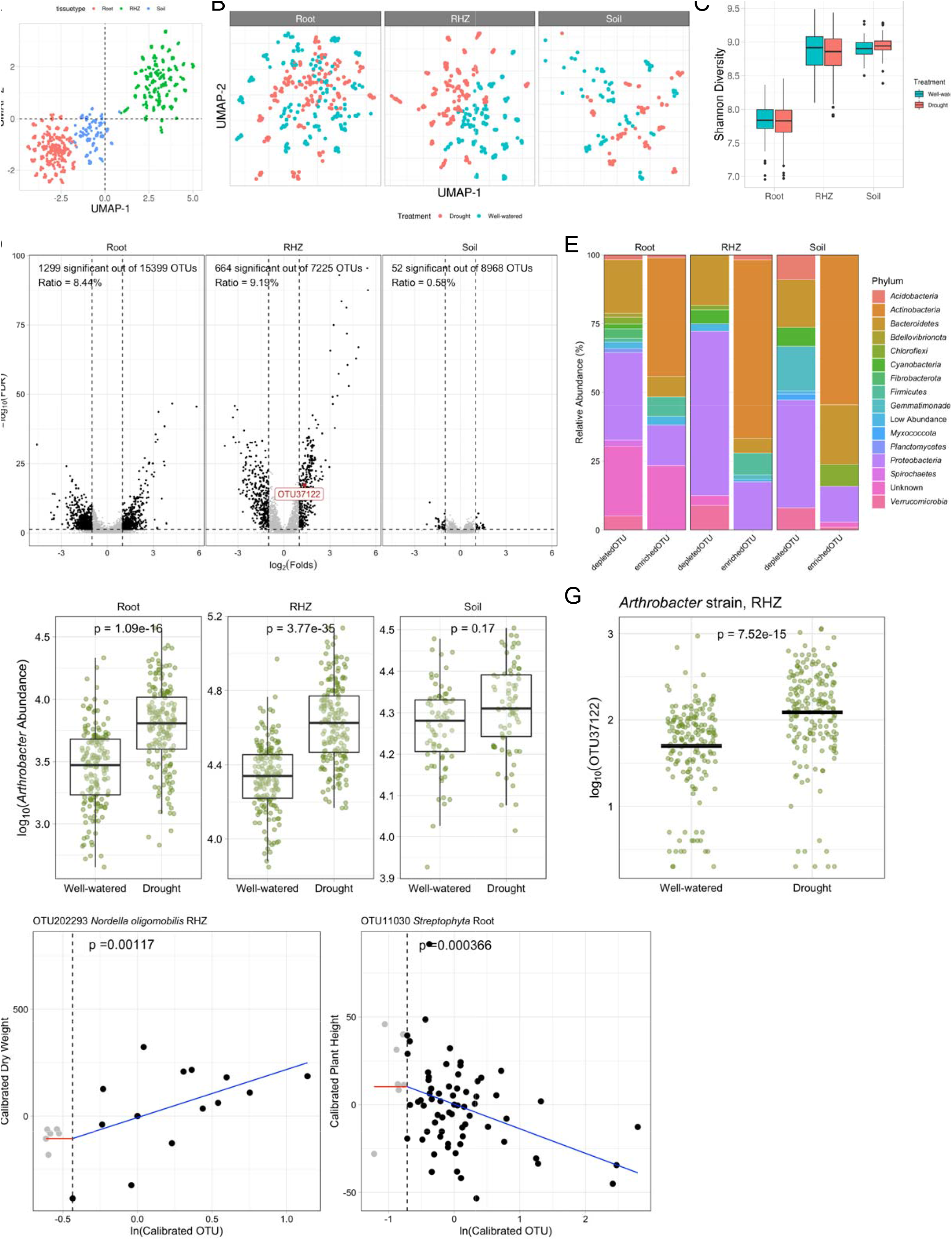
The sorghum microbiome with drought treatments in the field assay. A-B. The clustering of microbiome samples using supervised UMAP, with colors showing the tissue compartments (A) and the drought treatments (B). C. The Shannon diversity of sorghum microbiome samples. The solid dots represent the outliers and the colors show the drought treatments. D. For each tissue compartment (root, rhizosphere (RHZ) and soil), significantly differentially abundant OTUs between well-watered and drought are shown. The dashed lines show the cutoff thresholds: Folds < −2 or folds >2, FDR < 0.05. The dark red label shows OTU37122 *Arthrobacter* strain, which is enriched in RHZ under drought and negatively associated with plant growth phenotypes. E. Phylum-level distribution of the sorghum microbiota with significantly differentiated abundance (from panel D) within the tissue compartments under drought. F The abundance of *Arthrobacter* strains was enriched under drought in all three tissue compartments. ZINBGLMM models were fitted between the drought treatments with corrected p-values shown. G. The abundance of *Arthrobacter* strain OTU37122 was enriched under drought in rhizosphere. The horizontal bars represent medians. ZINBGLMM models were fitted between the drought treatments with corrected p-values shown. H. Examples of OTU abundance correlating with plant phenotype based on the change point hinge model (OTU202293: Positive association; OTU11030: Negative association). C and F, The horizontal bars within boxes represent medians. The tops and bottoms of the boxes represent the 75th and 25th percentiles, respectively. The upper and lower whiskers extend to data no more than 1.5× the interquartile range from the upper edge and lower edge of the box, respectively.

